# Tick bite-induced Alpha-Gal Syndrome and Immunologic Responses in an Alpha-Gal Deficient Murine Model

**DOI:** 10.1101/2023.11.09.566281

**Authors:** Surendra Raj Sharma, Shailesh K. Choudhary, Julia Vorobiov, Scott P. Commins, Shahid Karim

**Affiliations:** School of Biological, Environment and Earth Sciences, The University of Southern Mississippi, Hattiesburg, MS 39406, USA; Department of Medicine & Pediatrics, University of North Carolina, Chapel Hill, NC 27599-7280, USA

**Keywords:** alpha-gal, tick, *Amblyomma americanum*, alpha-gal knockout mice, delayed allergic responses, food allergy, mammalian meat, red meat allergy, hypersensitivity, Alpha-Gal Syndrome, *Amblyomma maculatum*

## Abstract

**Introduction:** Alpha-Gal Syndrome (AGS) is a delayed allergic reaction due to specific IgE antibodies targeting galactose-α-1,3-galactose (α-gal), a carbohydrate found in red meat. This condition has gained significant attention globally due to its increasing prevalence, with more than 450,000 cases estimated in the United States alone. Previous research has established a connection between AGS and tick bites, which sensitize individuals to α-gal antigens and elevate the levels of α-gal specific IgE. However, the precise mechanism by which tick bites influence the host’s immune system and contribute to the development of AGS remains poorly understood. This study investigates various factors related to ticks and the host associated with the development of AGS following a tick bite, using mice with a targeted disruption of alpha-1,3-galactosyltransferase (AGKO) as a model organism.

**Methods:** Lone-star tick (*Amblyomma americanum*) and gulf-coast tick (*Amblyomma maculatum*) nymphs were used to sensitize AGKO mice, followed by pork meat challenge. Tick bite site biopsies from sensitized and non-sensitized mice were subjected to mRNA gene expression analysis to assess the host immune response. Antibody responses in sensitized mice were also determined.

**Results:** Our results showed a significant increase in the titer of total IgE, IgG1, and α-gal IgG1 antibodies in the lone-star tick-sensitized AGKO mice compared to the gulf-coast tick-sensitized mice. Pork challenge in *Am. americanum* -sensitized mice led to a decline in body temperature after the meat challenge. Gene expression analysis revealed that *Am. americanum* bites direct mouse immunity toward Th2 and facilitate host sensitization to the α-gal antigen, while *Am. maculatum* did not.

**Conclusion:** This study supports the hypothesis that specific tick species may increase the risk of developing α-gal-specific IgE and hypersensitivity reactions or AGS, thereby providing opportunities for future research on the mechanistic role of tick and host-related factors in AGS development.

## Introduction

Alpha-gal syndrome (AGS) is an atypical allergic reaction to galactose-α-1,3-galactose (α-gal), a glycan present in all mammals except for catarrhine primates (Commins et al., 2009; Macher et al., 2008). Deactivation of the α-1,3-galactosyl transferase (α-1,3GT) gene in an ancestral Old-World species explains why humans, unlike other mammals, lack α-gal (Macher et al., 2008). As a result, the α-gal moiety becomes clinically significant because it triggers the production of anti-Gal antibodies in humans, including immunoglobulin M, A, and G (Macher et al., 2008; Galili, 1999). AGS, in contrast, is caused by a specific immunoglobulin E (sIgE) antibody response in sensitized hosts directed against α-gal. It usually leads to allergic reactions 2-6 hours after consuming red meat or its derivatives (Commins et al., 2009; Commins et al., 2014; Fischer et al., 2014).. The synthesis of α-gal-containing glycoconjugates involves a diverse family of glycosyltransferase enzymes (Berg et al., 2014; Roseman, 2001). Interestingly, these enzymes and the glycoconjugates they produce are mainly present in the cells, tissues, and fluids of mammals, excluding humans, apes, and old-world monkeys (Apostolovic et al., 2014; Galili & Avila, 1999; Hilger et al., 2016; Iweala et al., 2020; Takahashi et al., 2014). Consequently, the deactivation of α1,3GT in humans is believed to be the reason for developing an immune response to α-gal upon exposure to glycoconjugates containing α-gal antigens (Commins et al., 2014; Sharma and Karim, 2021).

Ticks are ectoparasites that can transmit various disease-causing pathogens, macromolecules, and other substances to humans (Adegoke et al., 2020; Bullard et al., 2019; Chmelař et al., 2016). Numerous scientific studies conducted globally have provided evidence that establishes a link between tick bites and the development of AGS (Araujo et al., 2016; Commins et al., 2011; Hamsten et al., 2013; Pacific & Van Nunen, 2015, Fisher et al., 2014). The rising prevalence of this emerging allergy has been observed in specific global regions, such as the United States (∼450,000 estimated cases (Thompson et al., 2023), where the increased tick population and their migration to new areas present a significant public health issue (Monzón et al., 2016; Raghavan et al., 2019). In certain major regions of the Southeastern U.S., it is estimated that up to 3% of the population has been affected by AGS, resulting in anaphylactic reactions (www.alphagalinformation.org, 2023). Furthermore, several other tick species worldwide, including *Ixodes holocyclus* in Australia, *Ixodes ricinus* and *Rhipicephalus bursa* in Europe, *Hyalomma marginatum* in Europe, *Haemaphysalis longicornis* in Japan, and *Amblyomma sculptum* in Brazil, have been identified as potential contributors to the development of AGS (Sharma and Karim, 2021).

The precise mechanism by which tick bites sensitize humans and contribute to the development of AGS is not fully understood. It is hypothesized that tick saliva, which contains α-gal antigens and salivary components, may trigger a host immune response and skew the immune system toward a TH2 response, resulting in the production of IgE antibodies that target α-gal (Araujo et al., 2016; Crispell et al., 2019; Choudhary et al., 2021). In fact, repeated tick bites have been observed to enhance the existing specific IgE antibody response (Commins et al., 2011; Kim et al., 2020; Hashizume et al., 2018). However, the relationship between glycosylated proteins containing α-gal in tick saliva and the process of α-gal sensitization or AGS induction in hosts requires further investigation, as these salivary factors may not be the sole determinant. It is worth noting that N-glycome profiling and proteome analysis have demonstrated the α-gal antigen in both salivary gland extracts and saliva of the lone-star tick (Am. americanum) and the black-legged tick (*Ix. scapularis*), while it is absent in *Am*. *maculatum* (Crispell et al., 2019). Indeed, previous research has demonstrated exposure to *Am. americanum* salivary gland extracts can induce the development of AGS in an AGKO mouse model (Choudhary et al., 2021). Recently, a case-control study provided evidence of an 11-fold increased risk of AGS in human hosts reporting tick bites (Kersh GJ, et al. 2023). Nevertheless, the specific conditions under which ticks, or other exposures trigger sIgE antibody production against α-gal, resulting in AGS, remain unclear. Consequently, it is essential to further investigate the tick and host-related factors associated with AGS induction after tick bites. This study aims to explore the role of tick bites in AGS development using AGKO mice and nymphal ticks, as well as examine how tick bites influence the host’s immune response and contribute to AGS development.

## Materials and Methods

### Ethics statement

All animal studies were conducted in strict accordance with the recommendations in the Guide for the Care and Use of Laboratory Animals of the National Institutes of Health, USA. The protocols for tick blood-feeding on mice and sheep were approved by the Institutional Animal Care and Use Committee (IACUC) of the University of Southern Mississippi (protocol #15101501.2, #19041801.2). All efforts were made to minimize animal distress and ensure their well-being throughout the procedures.

### Ticks and other animals

The lone-star tick (*Amblyomma americanum), hereafter* Aa or LST, and Gulf-Coast tick (*Amblyomma maculatum), hereafter* Am or GST, were maintained at the University of Southern Mississippi according to established methods (Patrick and Hair, 1975). Unfed adult lone-star ticks (Am*blyomma americanum*) were purchased from Oklahoma State University’s tick-rearing facility (Stillwater, OK, USA) and maintained at the University of Southern Mississippi using the previously described method (Patrick & Hair, 1975). Ticks were kept at room temperature at approximately 90% humidity with a photoperiod of 14 hours of light and 10 hours of darkness before infestation on mice. The nymph ticks were fed on mice for biopsy tissue collection, depending on the experimental plan.

### Mice

Alpha-1,3-galactosyltransferase knockout (AGKO) mice on C57BL/6 background were obtained from Dr. Anthony D’Apice (Tearle et al., 1996). AGKO mice were bred and maintained in pathogen-free rooms under protocols approved by the University of Southern Mississippi, Animal Care and Use Committee (IACUC). Euthanasia was performed by anesthetizing animals with an intraperitoneal injection of 1.25% avertin (125-250 mg/kg body weight) followed by cervical dislocation.

### Mice sensitization and food challenge

Eight- to ten-week-old AGKO mice were used for the sensitization experiment. During the sensitization experiment, mice were anesthetized by intraperitoneal injection of a 10 mg/kg ketamine/xylazine mixture, and 15 nymphal ticks (*Am. americanum* and *Am. maculatum*) were infested on the mice ear days 0, 7, 21, 28 (Fig. 1). Ticks were permitted to attach on the mice before being housed in individual cages with wire platforms above water to capture the engorged ticks. For repeat tick exposures in mice, ticks were allowed to feed till repletion and rested for an additional three-day post-drop-off before the next challenge. Mouse blood was collected on days 3, 10, 24, and 31 to quantitate total IgE, IgG, α-gal IgG, and α-gal sIgE. Mice sensitized to α-gal and control mice were orally challenged with 400 mg of cooked pork kidney homogenate (PKH) in phosphate-buffered saline. Core body temperature was measured with a rectal probe (Fisher Scientific, USA), before and after the meat challenge every 15 minutes for 2 hours. Mice were subjected to repeated rectal probe insertion and were conditioned before the food challenge to mitigate temperature variation induced by the insertion of the rectal probe.

**Figure 1.**
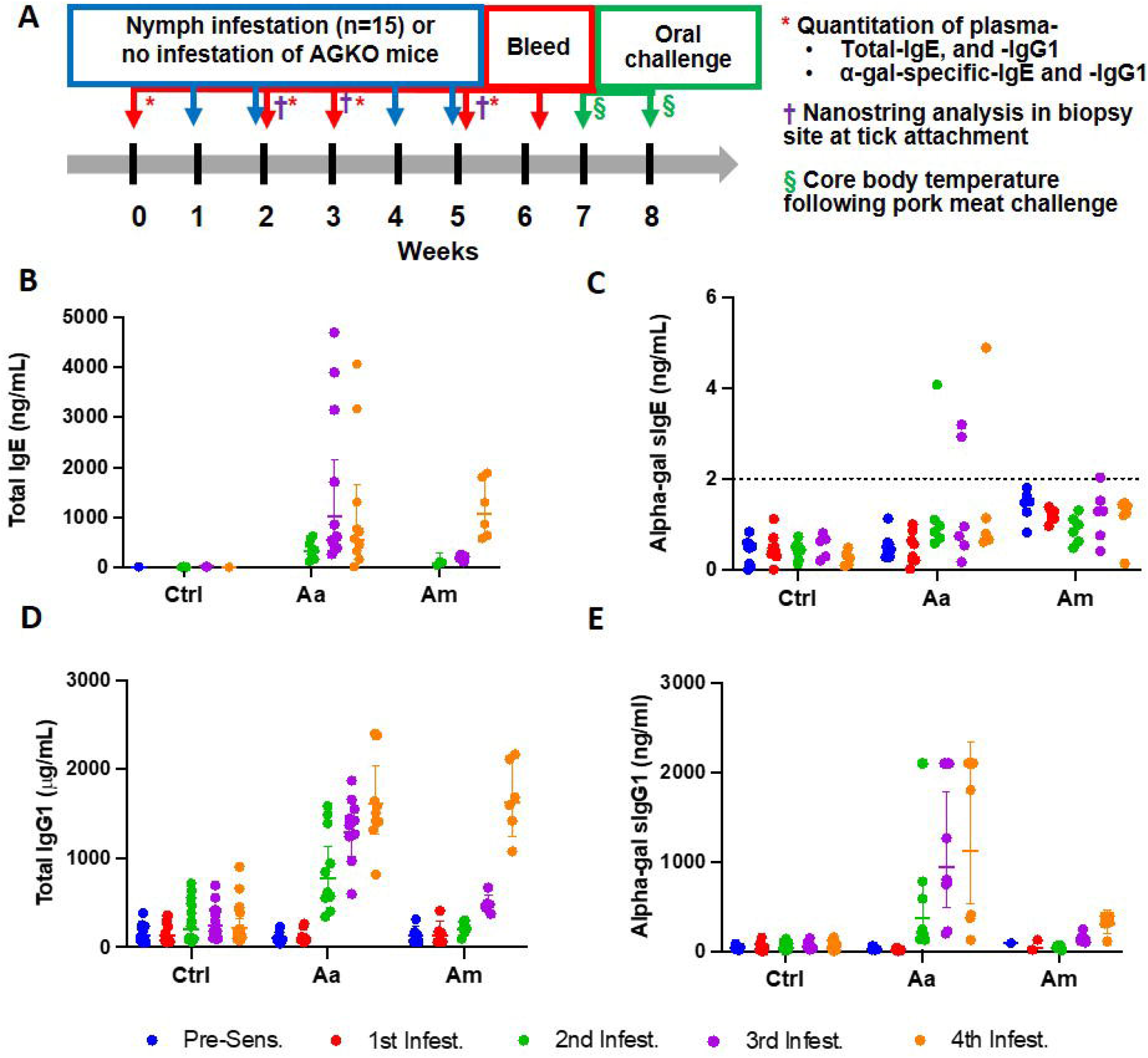
Sensitization of AGKO mice with tick nymphs. Schematic of alpha-gal sensitization, collection of biopsy site for immune responses, and oral meat challenge (A). Quantitation of total IgE (B), alpha-gal specific IgE (C), total IgG1 (D), and alpha-gal specific IgG1. Each dot represents an individual mouse and error bars represent the geometric mean with a 95% confidence interval. The dotted line shows the limit of detection of alpha-gal sIgE. Aa represents a group of mice (N=10) infested with *Amblyomma americanum*, Am represents a group of mice (N=6) infested with *Amblyomma maculatum* and Ctrl represents a group of mice with no infestation (N=16). Fifteen nymphs were used per mouse infestation.

### Quantitation of specific immunoglobulins

IgE was quantitated using the IgE Max Standard Set from Biolegend (San Diego, CA) according to the manufacturer’s instructions. Nunc Maxisorp plates were coated with 1X capture antibody or cetuximab (20 µg/ml) in carbonate-bicarbonate coating buffer to quantitate total IgE and αl-gal sIgE respectively. Briefly, plates were coated overnight at 4 °C, received four washes with PBS containing 0.05% Tween 20 (PBST; Sigma-Aldrich) and were blocked with 1% BSA in PBST for 90 minutes (min). Plasma samples (1:60 dilution for total IgE, 1:2 dilution for α-gal specific IgE) or standard were added to the plate and incubated for 2 hours (h) at room temperature (RT). Samples were incubated with 1X detection antibody for 1 h and avidin-HRP at RT in the dark for 30 min. 3,3’,5,5’-Tetramethylbenzidine (TMB) Peroxidase Substrate and Stop Solution (KPL, Gaithersburg, MD) was used to develop an enzymatic colored reaction. Plates were read on an Epoch Microplate Spectrophotometer (BioTek Instruments, Winooski, VT) and analyzed using Gen5 software. To quantitate IgG1 and α-gal sIgG1, plates were coated with capture antibody (goat anti-mouse IgG1, 1 μg/ml, SouthernBiotech) or cetuximab (20 µg/ml) respectively in carbonate-bicarbonate coating buffer overnight at 4 °C. Plates received four washes with PBST and were blocked with 3% fetal bovine serum (FBS) in PBST. Plasma samples (1:20,000 dilution in PBS containing 1% FBS for total IgG, 1:10 dilution for α-gal specific IgG) were incubated for 90 min at RT, and ELISAs were detected with HRP-conjugated goat-anti-mouse IgG1-HRP (Southern Biotech). To develop the enzymatic reaction TMB was used as described above. Antibody titer data further were analyzed using GraphPad Prism 9 (La Jolla CA). The two-way ANOVA with a mixed-model analysis and Tukey’s multiple comparison test was performed for statistical significance involving more than two groups while Mann-Whitney test was performed for single comparison.

### Skin biopsies, RNA Extraction, NanoString immunological assays, and analysis

Mice were anesthetized by intraperitoneal injection of a 10 mg/kg ketamine/xylazine mixture. Biopsy samples using 3 mm biopsy punches (Miltex, USA) were collected at the tick bite sites (1 site/animal) and stored in RNAlater Stabilization Solution (ThermoFisher, Waltham, MA, USA). During the partial feeding phase of ticks (3 days after infestation of mice), biopsy samples were obtained at the 1st, 2nd, and 3rd sensitization stages. To extract RNA from the skin biopsies, the RNeasy Plus kit (specifically, the Quick RNA miniprep plus kit, Zymo, USA) was utilized and RNA was quantitated with Qubit RNA HS (High Sensitivity) assay kit (Invitrogen, Waltham, MA). For the NanoString nCounter assay, 100 ng of RNA in 5 μL per sample was used for the Gene Expression Panel / CodeSet Hybridization protocol and each cartridge was hybridized for 18 hours on a thermal cycler set to 65℃ with a lid heated to 70℃. Once hybridization was complete, cartridges were moved to a nCounter Prep station and processed using the High Sensitivity protocol and then analyzed on the nCounter Pro Analysis System (NanoString Technologies, Seattle, WA, USA). NanoString Data was further analyzed by ROSALIND® (https://rosalind.bio/), with a HyperScale architecture developed by ROSALIND, Inc. (San Diego, CA). Read Distribution percentages, violin plots, identity heatmaps, and sample MDS plots were generated as part of the QC step. Normalization, fold changes, and p-values were calculated using criteria provided by Nanostring. ROSALIND® follows the nCounter® Advanced Analysis protocol of dividing counts within a lane by the geometric mean of the normalizer probes from the same lane. Housekeeping probes to be used for normalization are selected based on the geNorm algorithm as implemented in the NormqPCR R library1. An abundance of various cell populations is calculated on ROSALIND using the Nanostring Cell Type Profiling Module. ROSALIND performs a filtering of Cell Type Profiling results to include results that have scores with a p-Value greater than or equal to 0.05. Fold changes and pValues are calculated using the fast method as described in the nCounter® Advanced Analysis 2.0 User Manual. P-value adjustment is performed using the Benjamini-Hochberg method of estimating false discovery rates (FDR).

## Results

### Bites of Aa but not Am nymphs cause a robust production of IgE, IgG1 and α-gal-directed sIgG1 in an Alpha-gal KO mice mouse model

Pruritic reactions at the site of tick bites correlate with increased production of α-gal sIgE in humans, implying that sensitization to tick bites containing α-gal in saliva may require repeated tick bites (Hashizume et al. 2018; Wilson et al. 2021). We used nymphs of two tick species-Aa, known to secrete α-gal containing epitopes in the saliva, and Am, deficient in epitopes containing alpha-gal. Tick bite increased total and specific IgE and IgG1 production following sensitization (N=15 ticks infested, in an average total 7, ticks fed to repletion and dropped off per infestation) in AGKO mice (**Fig. 1A**). In control mice with no nymph infestation, total IgE was not detected in several mice at 60-fold dilutions of plasma. We, therefore, pooled the plasma samples across all four control infestations and measured total IgE (geometric mean (GM) = 7.44 ng/mL (n=6) with a 95% confidence interval (CI) of 3.44 to 16.06 ng/mL; **Fig. 1B**, **Table 1**). In mice with Aa and Am nymphs infested, total IgE increased significantly following the second infestation compared to the control. Notably, the increase in total IgE was 4.2-fold higher in Aa-infested mice than in Am-infested mice (p<0.05) and peaked following the third infestation with Aa (GM = 1022.53 ng/mL, CI 483.45 to 2162.75) and was significantly higher than Am-infested mice (GM = 194.16, CI 145.61 to 258.90). No significant difference in total IgE between Aa- and Am-infested mice was noted following the fourth infestation. We further detected α-gal specific IgE (sIgE) in two out of ten Aa-infested mice (**Fig. 1C**). In contrast, none of the Am-infested mice produced α-gal sIgE. An increase in IgG1 is associated with Th2 polarization and anaphylaxis in mice (Snapper and Paul 1987, Finkelman 2007). We, therefore, quantitated IgG1 and observed increased titer following tick infestation in both Aa- and Am-infested mice (**Fig 1D**, **Table 1**). Total IgG1 concentration following second and third infestations was significantly higher in Aa-infested mice (GM = 769.57, CI 522.06-1134.41 and GM = 1286.43, CI 1021.30-1620.40) in comparison to both Am-infested (GM = 200.94, CI 129.67-311.37 and 484.3, CI 397.86-589.53) and control (GM = 201.19, CI 126.38-320.28 and GM = 243.81, CI 170.48-348.69) mice. In contrast to the Aa-infested mice, the increase in total IgG1 titer remained low in Am-infested mice until the fourth infestation compared to control cohorts. Notably, α-gal specific IgG1 (sIgG1) was 13.1-fold, 6.5-fold, and 3.7-fold higher in Aa-infested mice than Am-infested mice following second, third, and fourth infestation, respectively; and these differences were significant (**Fig. 1E**, **Table 1**).

**Table 1.** Quantitaion of Immunoglobulines in control and tick-infested alpha-gal KO mice.

| Immunoglobulins | Treatments | Pre-sensitization |  | Tick infestation |  |  |  |  |  |  |  | Significance at 1st, 2nd 3rd and 4th infestation |
| --- | --- | --- | --- | --- | --- | --- | --- | --- | --- | --- | --- | --- |
|  |  |  |  | 1st |  | 2nd |  | 3rd |  | 4th |  |  |
|  |  | GM (GSD) | CI | GM (GSD) | CI | GM (GSD) | CI | GM (GSD) | CI | GM (GSD) | CI |  |
| IgE (ng/mL) | Control | †7.44 (2.08) | (3.44, 16.06) | †7.44 (2.08) | (3.44, 16.06) | †7.44 (2.08) | (3.44, 16.06) | †7.44 (2.08) | (3.44, 16.06) | †7.44 (2.08) | (3.44, 16.06) | Ctrl vs. Aa : -, ****, ****, ****<br>Ctrl vs. Am : -, **, ****, ****<br>Aa vs. Am : -, *, * *, ns |
|  | Aa |  |  |  |  | 317.43 (1.79) | (195.09, 516.48) | 1022.53 (2.85) | (483.45, 2162.75) | 538.33 (4.86) | (173.78, 1667.64) |  |
|  | Am |  |  |  |  | 74.91 (1.70) | (20.06, 279.70) | 194.16 (1.31) | (145.61, 258.90) | 1061.48 (1.67) | (617.77, 1823.88) |  |
| IgG1 (µg/mL) | Control | 125.36 (1.89) | (89.29, 175.99) | 138.26 (1.2) | (94.27, 202.77) | 201.19 (2.39) | (126.38, 320.28) | 243.81 (1.91) | (170.48, 348.69) | 212.96 (2.13) | (140.20, 323.50) | Ctrl vs. Aa : ns, ****, ****, ****<br>Ctrl vs. Am : ns, ns, ns, ****<br>Aa vs. Am : ns, ***, **, ns |
|  | Aa | 98.38 (1.58) | (70.87, 136.55) | 107.44 (1.64) | (75.50, 152.90) | 769.57 (1.72) | (522.06, 1134.41) | 1286.43 (1.38) | (1021.30, 1620.40) | 1606.93 (1.39) | (1265.69, 2040.20) |  |
|  | Am | 126.2 (1.85) | (66.07, 241.03) | 137.1 (2.05) | (64.61, 290.92) | 200.94 (1.52) | (129.67, 311.37) | 484.3 (1.21) | (397.86, 589.53) | 1630.5 (1.30) | (1240.52, 2143.08) |  |
| sIgG1 | Control | 34.79 (1.72) | (23.63, 51.22) | 26.92 (3.64) | (11.31, 64.09) | 43.03 (2.66) | (23.12, 80.07) | 53.76 (1.91) | (33.82, 85.47) | 48.53 (3.43) | (22.19, 106.16) | Ctrl vs. Aa : ns, ***, ****, ****<br>Ctrl vs. Am : ns, ns, **, ***<br>Aa vs. Am ns, **, ***, ** |
|  | Aa | 27.9 (1.63) | (18.51, 42.05) | 14.74 (2.1) | (8.67, 25.06) | 366.25 (3.02) | (166.14, 807.36) | 932.87 (2.45) | (490.84, 1772.98) | 1115.6 (2.80) | (534.00, 2330.63) |  |
|  | Am | 92.3 (1.0) |  | 44.34 (4.38) | (?, ?) | 35.19 (2.28) | (12.64, 97.93) | 142.85(1.37) | (101.92, 200.21) | 301.06 (1.56) | (199.0, 455.47) |  |

GM, geometric mean; GSD, geometric standard deviation (GSD); CI, 95% confidence intervals; ns, not significant, \*, p<0.05; \*\*, p<0.01; \*\*\*, p<0.001; \*\*\*\*, p<0.0001
† Toal IgE detected above background were pooled acrossse all time points

### Tick bite-sensitized mice exhibit a drop in core body temperature after the red meat challenge

We have previously reported that oral challenge of sensitized mice with pork kidney causes a more consistent and faster reaction (less than 2 hours) due to the high content of α-gal in heavily glycosylated proteins such as angiotensin I-converting enzyme (ACE I) and aminopeptidase N (AP-N) found in pork kidney (Choudhary et al., 2020). We, therefore, challenged tick bite-sensitized AGKO mice with 400 mg of cooked pork kidney homogenate (PKH) orally to study the allergic response to α-gal. Body temperature decline was measured every 15 minutes over two hours with a rectal thermometer. A drop of mean body temperature between 1.5-3.0°C was taken as a sign of mild anaphylaxis and below 3.0°C as severe anaphylaxis. In Aa-infested mice, severe anaphylaxis was noted at 30 min after PKH with a mean temperature decline of 5.8°C. It was significantly different than control mice or *Am*-infested mice (**Fig. 2**). The body temperature reached its nadir 60 minutes after the challenge, and the mice showed symptoms of reduced activity and labored breathing (data not shown). In contrast, no significant decline in body temperature was observed in control and Am-infested mice, where a drop in the body temperature was less than 1.5°C following the PKH challenge. Our results suggest that infestation with Aa but not Am causes anaphylaxis in AGKO mice.

**Figure 2.**
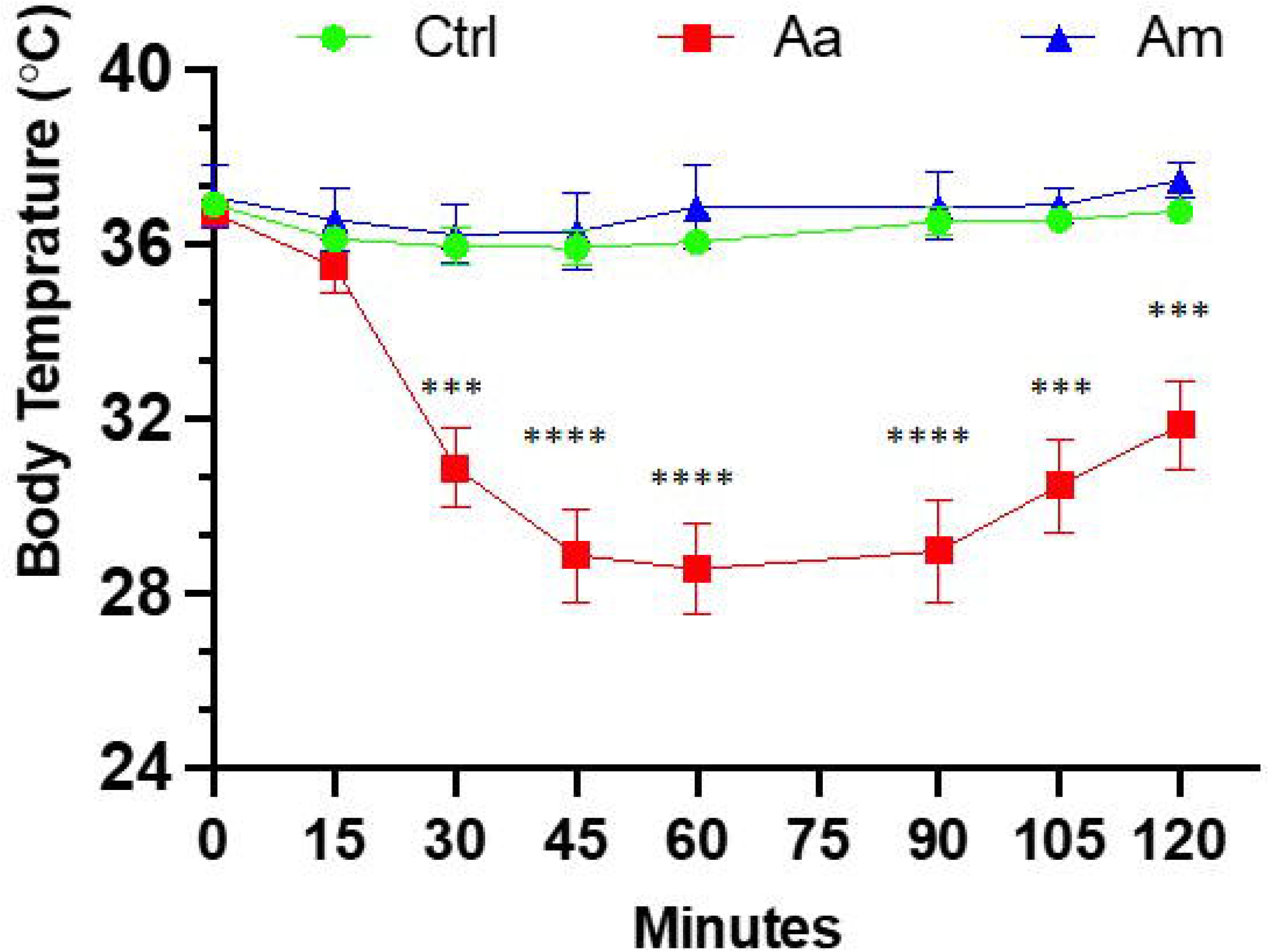
Anaphylaxis in tick infested mice following post-oral challenge with 400 mg of cooked pork kidney homogenates. Body temperature was recorded at baseline and post-oral challenge and a drop in body temperature >3 was considered as a sign of severe anaphylaxis. The two-way ANOVA with Tukey’s multiple comparison test was performed for statistical significance; *** P<0.001; **** P<0.0001. Aa, *Amblyomma americanum;* Am, *Amblyomma maculatum;* Ctrl, no infestation.

### AGKO mice sensitization using repeat tick infestation and nymph engorgement rate

To investigate whether repeated tick exposure in AGKO mice leads to acquired tick resistance (ATR) and host rejection, we conducted an experiment involving sensitizing ticks to AGKO mice through four repeated infestations. The engorgement data indicated that repeated exposure to Aa ticks resulted in a small but significant decline in nymph body weight following the second infestation. The Aa nymph body weight decline peaked following the third infestation; no further reduction was noted at the fourth infestation (**Fig. 3 A**). In contrast, no significant impact on tick engorgement was observed in Am nymphs (**Fig 3B**).

**Figure 3:**
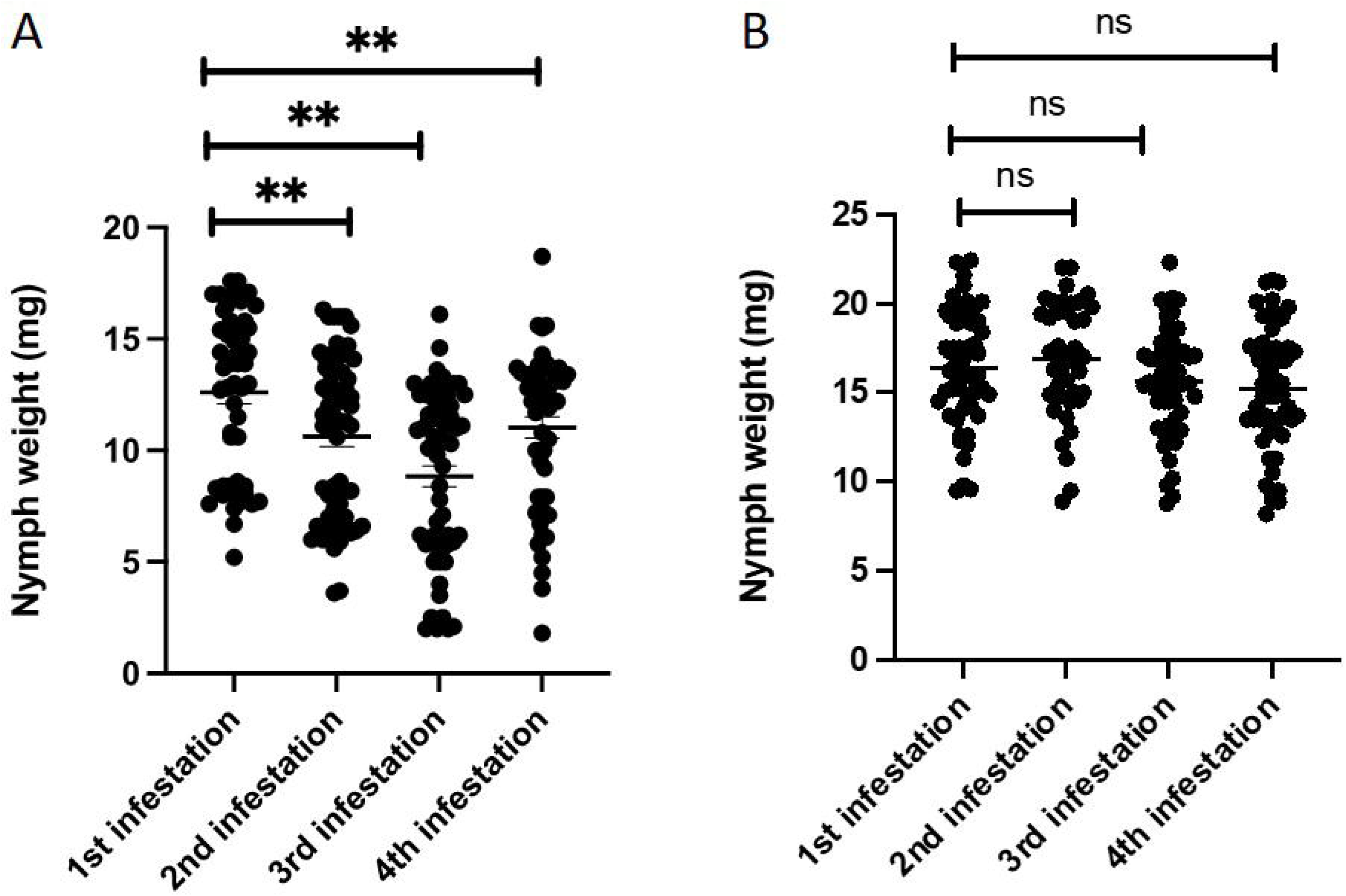
Repeated sensitization of AGKO mice via nymph tick bite and analysis of tick engorgement. (A) Engorgement weight of *Amblyomma americanum* nymph tick recovered during different stage of sensitization of AGKO mice (N=5). (B) Engorgement weight of *Amblyomma* maculatum nymph tick recovered during different stage of sensitization of AGTKO mice (N=5). Statistical test-Student t test; ns: non-significant, ** P<0.05.

### Differential gene expression at tick bite site

Skin biopsy was performed at the site of Aa tick attachment; RNA was extracted and quantitated as described in the Methods section. The Mouse Host Response Panel supplemented with primers for Bach2, Clec7a, Ighg1, Mmp13, Mmp8, Rorc, S100a, Sp140, TSLP, and Ybx3, was used for expression profiling of 783 genes via the digital multiplexed NanoString™ nCounter analysis system. Data were analyzed using the Rosalind NanoString™ Gene Expression platform. A sample correlation heatmap was drawn in which the data matrix contains correlation values between samples, with the darkest blue representing the strongest correlation (**Fig 4A**). Replicates of tick-infested mice strongly correlated; however, some correlation was also observed between control and 3/18 tick-infested samples. Fold change and p-values were calculated using the fast method, and p-value adjustment was performed using the Benjamini-Hochberg method of estimating false discovery rates (FDR). Setting a filter at fold change to ≥ 1.5 and ≤ -1.5 and p-adj to 0.05, first-, second-, and third-tick infestation resulted in gene expression change of 283 (162 upregulated, 111 downregulated), 158 (122 upregulated, 36 downregulated, and 313 (248 upregulated, 65 downregulated) genes respectively. When data were compared for all three tick infestations versus control, 329 genes were differentially expressed, of which 237 were upregulated, and 92 were downregulated. Volcano plot and heat map (**Fig. 4B-C**) show that data with a fold change of ≥2.0 and ≤ -2.0 and p-adj to 0.01 that identified 169 upregulated and 53 downregulated genes.

**Figure 4.**
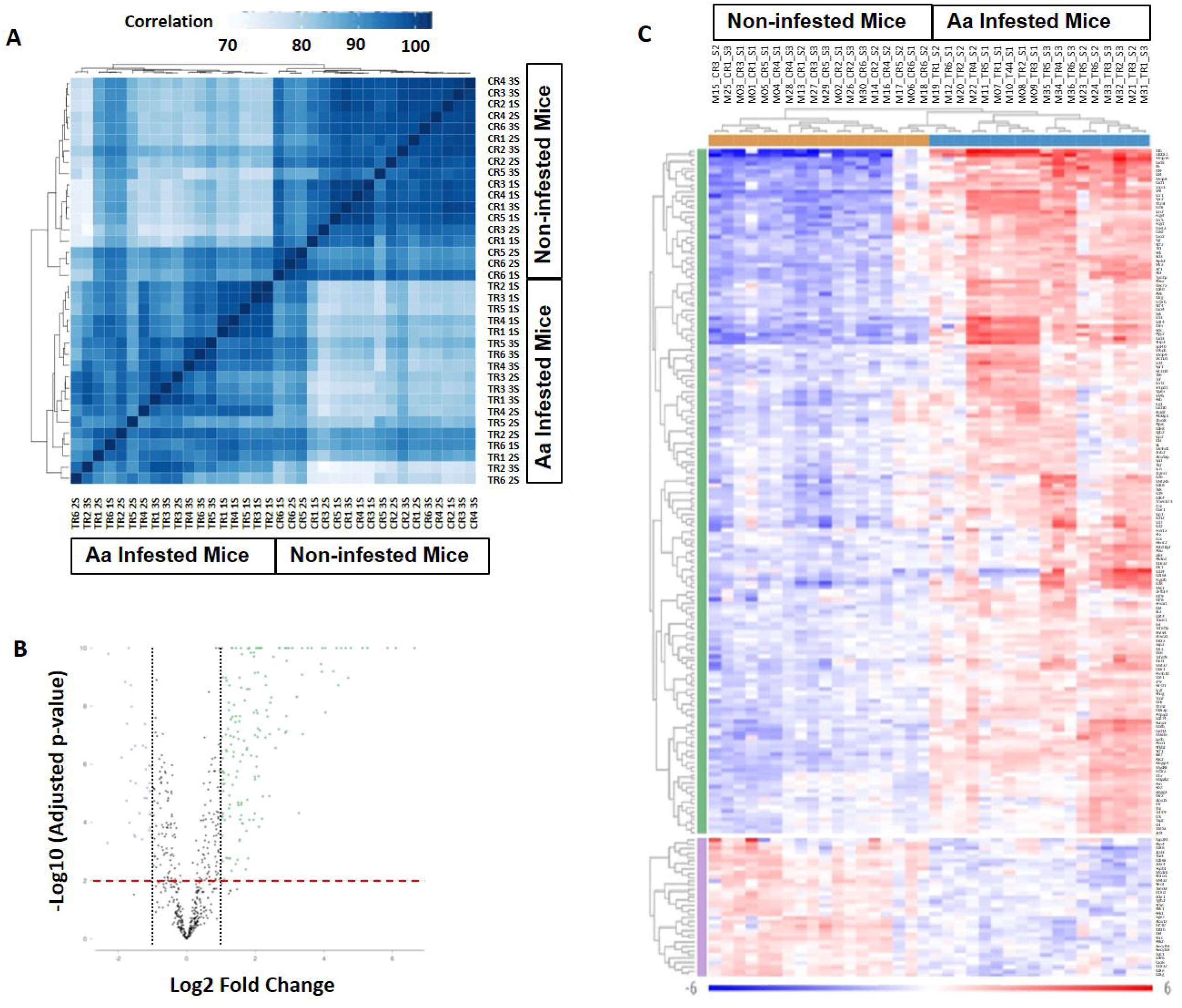
Differential gene trascription in biopsy site at tick attachment. (**A**) Sample correlation heatmap. (**B**) Volcano plot with dotted line displays mean gene expression fold change ≥ 2.0 and ≤ -2.0 between tick-infested mice and control infestation. The dashed red line shows where adjusted p value = 0.01. (**C**) Heat map of genes displaying two-fold change and p-Adjusted value ≥0.01. The legend display mapping to row-wise Z-scores. Aa, *Amblyomma americanum*; 1S, first sensitization; 2S, second sensitization; 3S, third sensitization; TR1-TR6, Aa infested mice; CR1-CR6, non-infested mice.

### Regulation of chemokine, cell adhesion, vascular permeability, and leucocyte migration

Cell type profiling was based on the NanoString™ Cell Type Profiling Module and filtered using Rosalind to include data with a p-value < 0.05 demonstrating that monocytes and NK56^dim^ cells were predominantly detected at the site of tick attachment in comparison to no tick infestation (**Fig. 5A**). In keeping with this, we observed an upregulation of monocyte chemokine genes *Ccl3* and *Ccl4* (**Fig. 5B**). The induction of *Ccl3* (35-fold) and *Ccl4* (21-fold) peaked following the 1st tick attachment with a concomitant increase in their receptor, *CCR1*. Other CC chemokines (*Ccl2*, *Ccl7*, *Ccl8*, *Ccl12*, *Ccl24*) were also robustly induced; however, expression of these peaked following additional tick infestations. The induction of *Cxcl1*(12-fold), *Cxcl3* (64-fold), and *Cxcl10* (6-fold) peaked following the 1st tick infestation, while *Cxcl5* expression rose 39-fold following the 3rd tick infestation. Gene sets associated with tissue damage and inflammation were induced in tick-attached mice. *Sell*, which encodes the adhesion molecule L-selection, was highly upregulated following tick infestation (**Fig. 5C**). Plasminogen activator *Plau* and plasminogen activator *Plaur*, and the matrix metallopeptidases *Mmp8*, *Mmp9*, and *Mmp13* were all induced several fold (**Fig. 5D**). Furthermore, we observed increased transcription of *Ptgs2*, a key enzyme in prostaglandin biosynthesis and associated receptor *Ptger4* and bradykinin receptor *Bdkrb1*.

**Figure 5.**
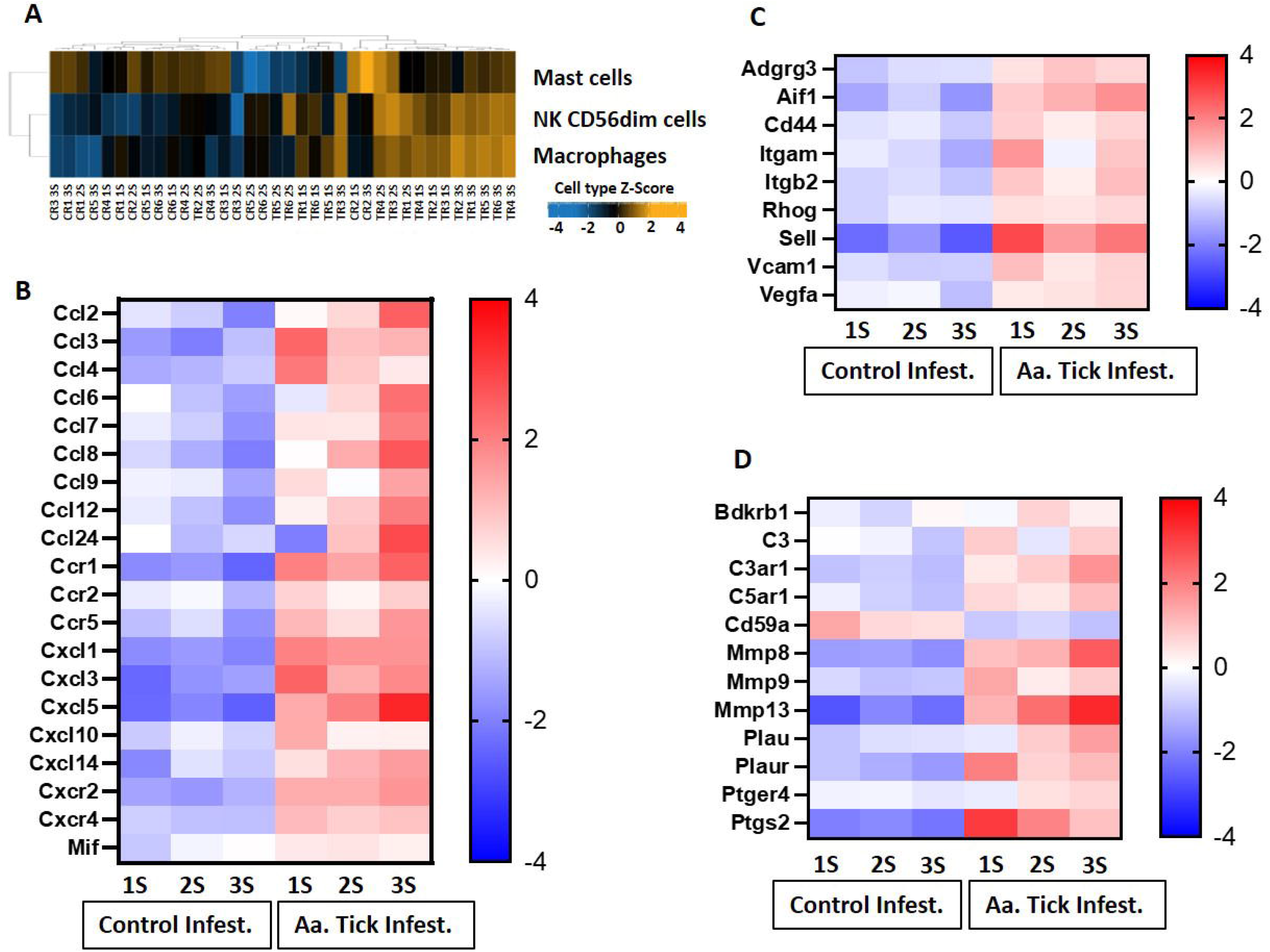
Heat map representation of differentially transcribed genes involved in cell migration and recruitment of leucocytes to tick-infestation sites. **(A)** Cell population present on site of tick attachment based on gene characteristics determined by NanoString cell type profiler. **(B)** Chemokine and chemokine receptors, **(C)** Cell adhesion, vascular permeability and leucocyte migration. **(D)** Plasminogen and complement activation. Aa, *Amblyomma americanum*; 1S, first sensitization; 2S, second sensitization; 3S, third sensitization.

### Induction of cytokines, interferons, inflammasome and other inflammatory mediator genes

Infestation of mice with ticks resulted in a massive induction of genes involved in IL-1β secretion and activation at the first timepoint (*IL-1β* increased 831-fold; **Fig. 6A**). Induction of *IL-1β* coincided with upregulation of pattern recognition receptors (PRR) of distinct classes which included the following: (i) Toll-like receptors (TLR) members *Tlr1*, *Tlr2*, *Tlr5*, *Tlr6* and *Tlr8* (ii) Nod-like receptor (NLR) member *Nod2* and C-type lectin receptor (CLR) member *Clec7a* (**Fig. 6B**). *CD14*, which is also a co-receptor for bacterial lipopolysaccharide (LPS) was induced 20-fold (Wright et al. 1990). *MyD88* transcripts were induced, and this critical member of TLR signaling is involved in both NF-kB and AP-1 activation (Kawasaki and Kawai 2014). Downstream transcription factors *Map2k2*, *Mapkapk2*, *Batf* and *Junb*, involved in the nuclear translocation of AP-1 and *Nfkb2*, *Nfkbia* and *Bcl3*, which activate NF-kB, were all induced (**Fig. 6C**). Furthermore, upregulation of *Tnfα* (six-fold), and its receptor *Tnfrsf1a* were noted, which could activate NF-κB downstream targets as well. Induction of pro-IL-1β by both NF-kB and MAPK pathways was supported by NOD [nucleotide oligomerization domain]-, LRR [leucine-rich repeat]-, and PYD [pyrin domain]-containing protein 3 (NLPR3) inflammasome activation; transcription of *Nlrp3* was induced 45-fold following the 1st infestation of mice with ticks (**Fig. 6B**) (Jo et al. 2016).

**Figure 6.**
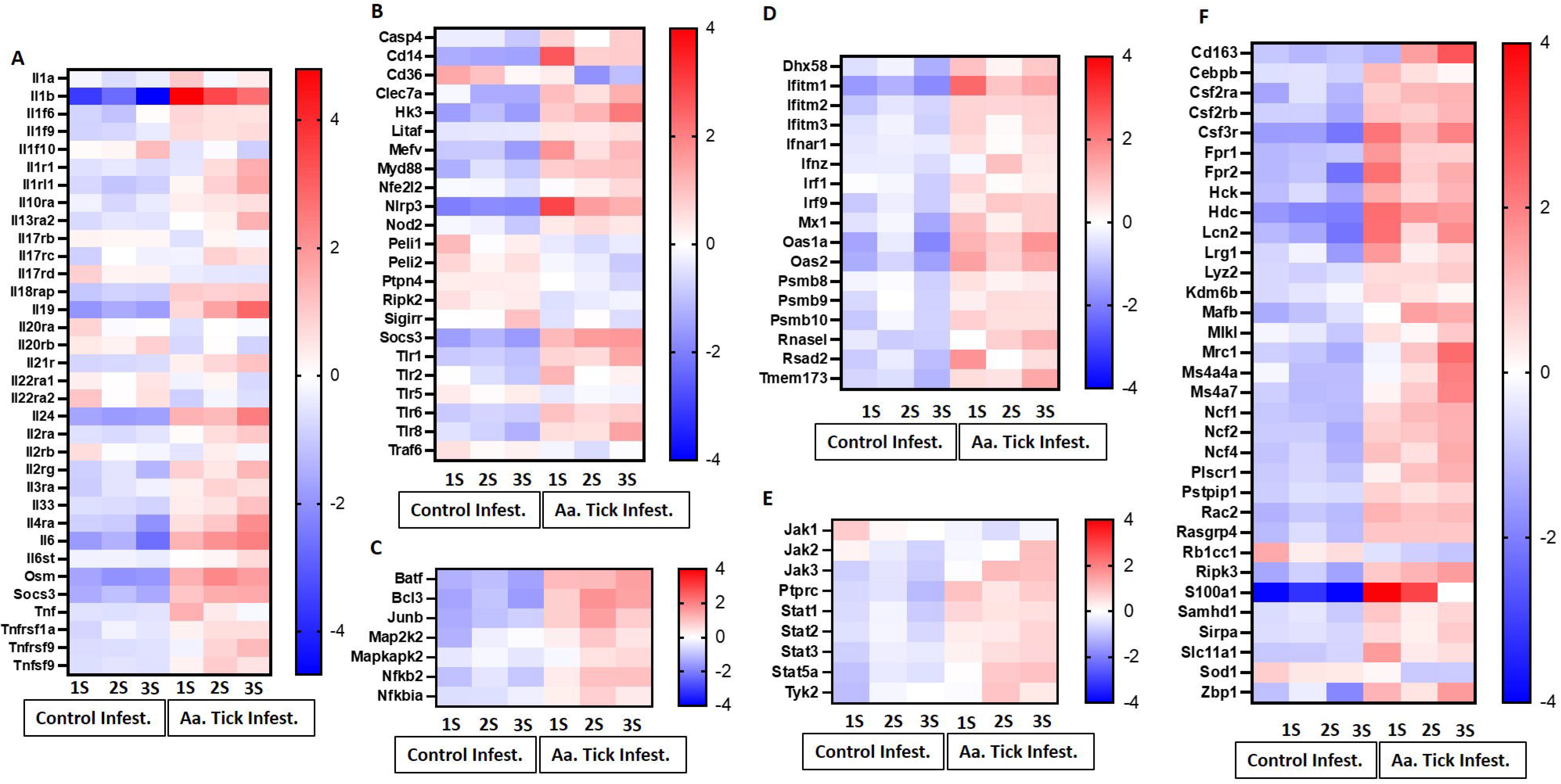
Heat map representation of differentially transcribed genes involved in inflammatory and anti-inflammatory responses. **(A)** Cytokines and cytokine receptors. **(B)** Pattern recognizing receptor (PRR), toll-like receptor (TLR) and inflammasome activation. **(C)** MAPK and NF-κB signaling pathway. **(D)** Interferon and antiviral responses. **(E)** JAK-STAT signaling pathway. (**F**) Other inflammatory response genes.

Several negative modulators of TLR signaling were induced, which would lead to inhibition of both the signaling complex formation of pro-IL-1β transcription as well as inflammasome inactivation (**Fig. 6B**). *Traf6*, which is involved in both MyD88-dependent and TRIF-dependent TLR signaling pathways was downregulated, whereas *Socs3*, which suppresses the activation of the MyD88-dependent pathway, was upregulated. The IL-20 subfamily of the IL-10 cytokine family elicits innate defense mechanisms from epithelial cells against extracellular pathogens (Rutz et al. 2014). Following the first tick infestation, two members of this subfamily *Il19* and *Il24* were induced and further increased following more tick infestations (**Fig. 6A**). *Il6* and *Osm*, which belongs to the IL-6 cytokine family, were induced following the first tick infestation and their transcription increased following subsequent tick infestation along with their receptor *Il6st* (**Fig. 6A**).

Consistent with the induction of PRR and TLR-mediated signaling, all three members of the CD225 superfamily of interferons: *Ifitm1*, *Iftim2*, and *Ifitm3* were upregulated (**Fig. 6D**) (Yanez et al. 2020). JAK/STAT signaling pathway is critical to cytokine-receptor signaling and induces key mediators of inflammations (Tangye et al. 2023). Tick infestation resulted in the upregulation of *Stat1*, *Stat2*, *Stat3* and *Stat5a* (**Fig. 6E**). Three members of the Janus kinase family *Jak2*, *Jak3* and Tyk2 were upregulated while *Jak1* was downregulated. During an inflammatory state, common β-chain cytokines such as GM-CSF, IL-3 and IL-5 further regulate the inflammatory response in a cell- and tissue-specific manner. We observed an increase in the transcription of *Csf2ra*, *Csf2rb* and *Csf3r* of *GM-CSF* and *G-CSF* which promote the generation of granulocytes and antigen-presenting cells (**Fig. 6F**). Neutrophils and monocytes are the first leucocytes to be recruited to an inflammatory site. Transcripts of *Fpr1* and *Fpr2*, the high-affinity receptors for N-formyl-methionyl peptides (fMLP), a powerful neutrophil chemotactic factor, were upregulated (Boulay et al. 1990).

### Antigen presentation and Th2 polarization

Following the second infestation of mice with ticks, indicators of a switch in the immune response from an innate to an adaptive response were present. *Cd6*, which expresses a costimulatory molecule to promote T-cell activation in response to PRR, was induced (**Fig. 7A**)(Sarrias et al. 2007). Several MHC class I transcripts, including *H2-D1*, *H2-K1*, *H2-M3*, *H2-Q10*, and *H2-T23*, were upregulated as well. *Rorc* was downregulated and, as a suppressor of IL-2 in mice, this decrease would lead to further T cell activation (He et al. 1998). Importantly, the transcription factor *Foxp3* was significantly induced following the 2nd infestation, consistent with a Treg response to suppress inflammation (**Fig. 7B**)(Ono et al. 2007). No significant change in gene expression was observed for Th1 transcription factor *Tbx21* (**Fig. 6C**) (Szabo et al. 2000). MHC class II molecules are involved in Th2 response, and *Cd209e*, which is involved in dendritic cell-mediated APC, was induced (**Fig. 7A**) (Angel et al. 2009). Further, the transcript for cathepsin, *Ctss*, involved in removing the invariant chain from MHC class II molecules and MHC class II antigen presentation was upregulated (Baranov et al. 2019). Several receptors and costimulatory molecules, such as *Icos*, *Cd40*, *Cd40lg*, and *Cd80* were induced for efficient antibody response to T-cell-dependent antigens (**Fig. 8A**). *Cd84*, which prolongs T-cell: B-cell contact, optimal T follicular helper function, and germinal center formation, was induced (Cannons et al. 2010). In keeping with the switch to an adaptive response, induction of *Il6* was observed that supports the differentiation of B cells into immunoglobulin-secreting cells as well as the development of T follicular helper (Tfh) cells (Dienz et al. 2009, Eto et al. 2011). Cytokines IL-4, IL-13, IL-21, and IL-33 are involved in different aspects of immunoglobulin secretion, such as Th2 differentiation, generation of Tfh cells, and formation of the germinal center (Coffman et al. 1986, Defrance et al. 1994, Schmitz et al. 2005, Dienz et al. 2009). We observed an increase in transcripts of the pro-Th2 cytokine Il33 and its receptors *Il1rl1*, and *Il4* and its receptors *Il4ra*, *Il21r*, and *Il13ra2* (**Fig. 8A-G**). Transcription of *Il21* was significantly induced following the 2nd tick infestation; however, no induction was observed for *Il13* (data not shown). *Maf*, which activates the expression of IL4 in Th2 cells, *Pik3ap1*, which contributes to PI3K-Akt-mediated BCR signaling, and *Pik3cd*, which mediates proliferative of B cells in response CD40 and IL-4 stimulation, were all induced (Ho et al. 1998, Troutman et al. 2012, Avery et al. 2018). Consistent with the induction of Th2 differentiation genes, increased expression of the transcripts of Fc receptors such as *Fcer1a*, *Fgr1*, *Fcgr2bm*, *Fcgr4*, and *Ms4a2* were observed. Furthermore, increased gene expression of *Alox15*, *Atf4*, and *Xbp1*, involved in ER stress response and immunoglobulin secretion, was observed (*So, 2018*). Our data suggest that Th2 differentiation and immunoglobulin production predominate the immune response following the 2nd tick infestation.

**Figure 7.**
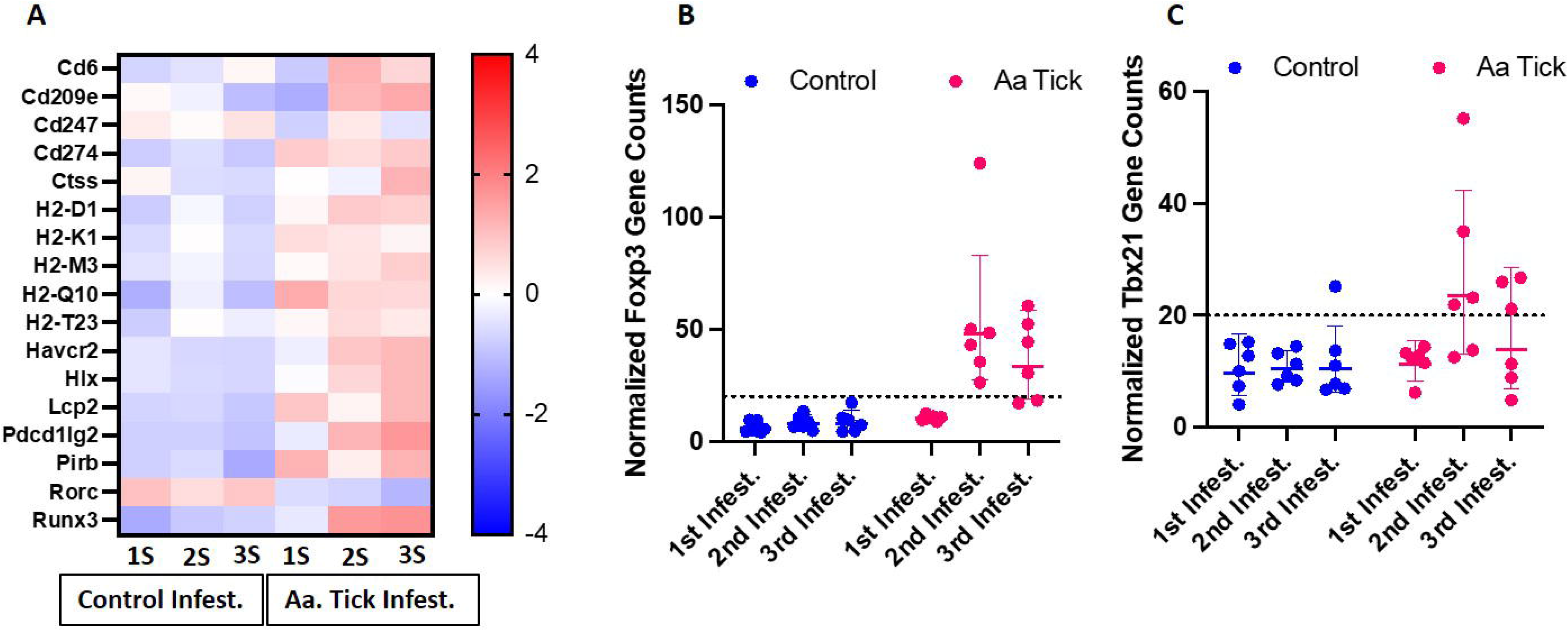
Modulation of genes involved in antigen presentation and Th1 response. **(A)** Heat map showing representation of differentially transcribed genes involved in antigen presentation (A). (B-C) Dot plots showing transcript counts of Treg transcription factor Foxp3 and Th1 transcription factor Tbx21. Each dot represents an individual mouse. Dotted line show the limit of detection.

**Figure 8.**
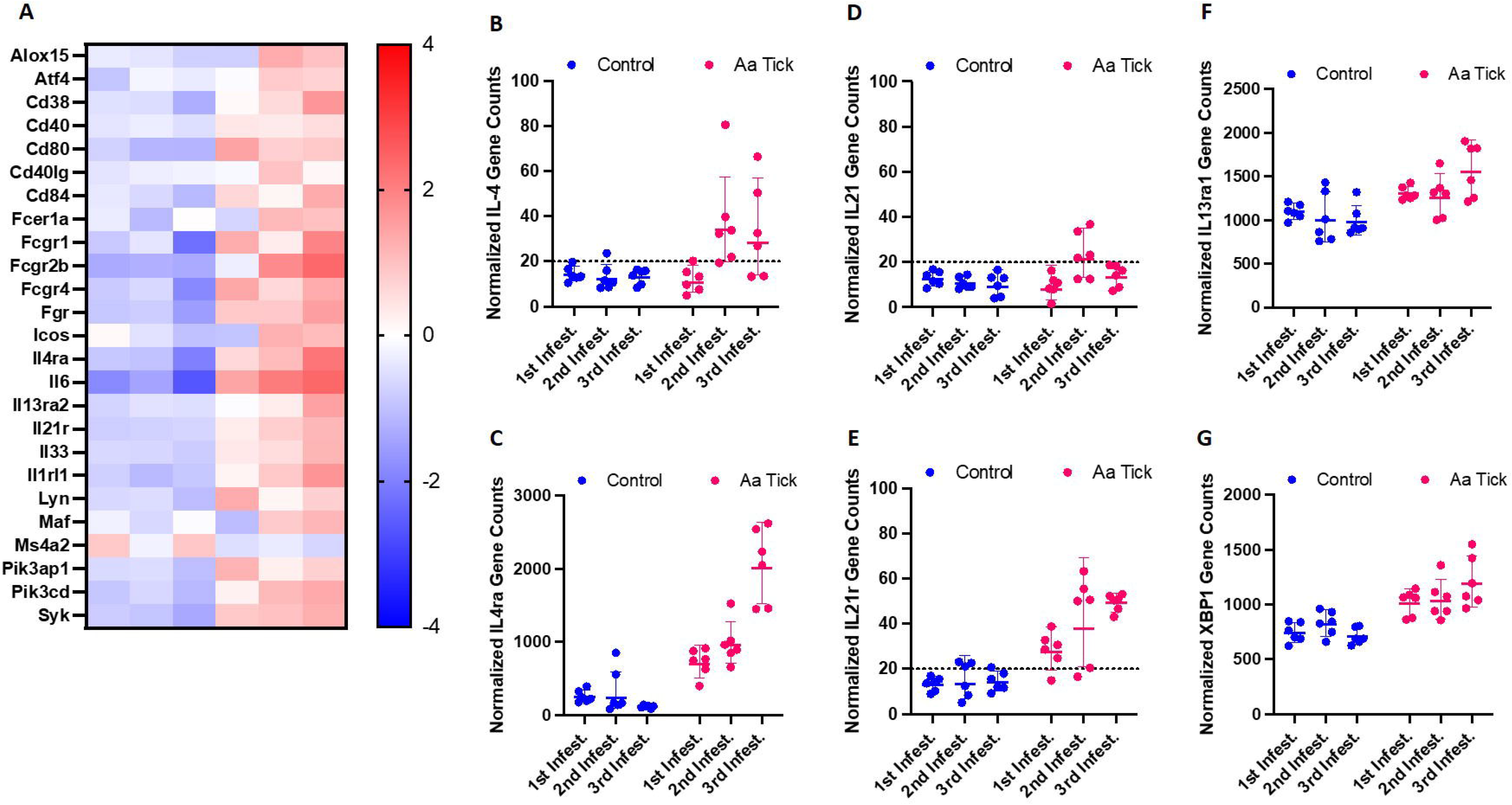
Modulation of genes involved in B cell signaling and Th2 response. Heat map showing representation of differentially transcribed genes involved in B cell signaling and Th2 response (A). Dot plot showing differential transcription of cytokine IL-4 and its receptor (B-C), cytokine IL-21 and its receptor (D-E), cytokine IL-13 receptor (F) and ER stress response factor XBB-1. Each dot represents an individual mouse. Dotted line show the limit of detection.

## Discussion

The current study reports the first small animal model that utilizes live, attached ticks to induce AGS as observed in humans. Through this, we tested the hypothesis that lone-star tick or Aa bite induces a Th2 immune response in the mammalian host, reorienting the immune system to produce IgE antibodies responsible for AGS. We compared immune-related gene expression profiles of the tick-sensitized and unsensitized AGKO murine samples to identify relevant pathways involved in the IgE response following tick bites. Based on AGS patient tick bite histories, tick species prevalence, in the regions with high AGS incidences, it is thought that α-gal sIgE develops in humans after being bitten by certain tick species. For instance, *Am. americanum* is associated with this response in the United States, while in Australia, Europe, Japan, and Brazil, *Ixodes holocyclus*, *Ixodes ricinus*, *Haemaphysalis longicornis*, and *Amblyomma sculptum*, respectively, have been implicated in AGS (Commins et al., 2011; Commins 2020; van Nunen et al., 2009; Crispell et al., 2019; Sharma and Karim, 2021).

Interestingly, epidemiologic surveillance studies have found a correlation between the geographic distribution of Aa ticks, high titer of α-gal sIgE antibody in AGS patients (Commins et al., 2014; Kersh et al., 2023). Beyond epidemiologic evidence from a recent case-control study, no direct and definitive evidence of tick bite induced AGS exists in humans. In keeping with this, the mechanisms by which *A*a tick bites induce high titer sIgE production and initiate delayed allergic response after exposure to red meat in humans are poorly understood. Among available allergy mouse models, we and others have previously utilized an AGKO mouse model, which mimics humans as an “alpha-gal-deficient host” and reported the induction of α-gal sIgE following immunization using partially fed salivary gland extracts and larval protein extracts with adjuvants, respectively (Choudhary et al., 2021; Chandrasekhar et al., 2019). These sensitization methods have their own set of limitations. For example, employing tick salivary extract or larval homogenate for sensitization permits the utilization of salivary protein extracts and offers evidence suggesting ticks’ recognition of their role in AGS development. Despite their utility, these methods fail to replicate the intricate natural process of tick attachment and the secretion of tick salivary factors during feeding, which is crucial for host sensitization.

Additionally, the salivary extract contains several proteins; many of those are non-secretory glycosylated proteins and may not be part of tick saliva but still can drive antibody response against the α-gal antigen in the host. Consequently, the host response driven by cutaneous injection may not truly reflect the sensitization process during prolonged and repeated tick feeding. Furthermore, no direct evidence demonstrates that mouse sensitization following repeated tick feeding causes AGS induction in the AGKO murine model. Therefore, we aimed to develop a tick sensitization model by sensitizing AGKO mice through repeated nymphal tick feeding. This model was used to study the host’s immune response to tick bites using tick species *Aa* and *Am* to determine their role in inducing the sIgE response. The alpha-gal signature in the saliva and salivary glands of the lone-star tick has been reported (Crispell et al., 2019; Park et al., 2020; Sharma et al., 2021), and subcutaneous injection of tick salivary glands extract has also been reported to induce high titer IgE response in AGKO murine model (Choudhary et al., 2021). Figure 1 shows a gradual increase in the total IgE and IgG1 levels in tick-sensitized mice compared to the control mice group.

Intriguingly, the total IgG titer of Am-sensitized mice was similar following 4^th^ sensitization however in the Aa sensitized mice, total IgG titer peak earlier (from 3^rd^ sensitization). It is expected to see an elevated IgG titer after any tick species bite; however, Am bite-induced high titer may correlate with the voracious and aggressive feeding behavior and high amount of salivary protein injected into the host compared to Aa. Noticeably, Aa-sensitized mice showed a significantly higher α-gal IgG1 antibody titer after 3^rd^ and 4^th^ infestation, indicating salivary antigens from Aa boosts anti-α-gal host responses (Figure 1B). The total IgE titer significantly increased after the 4^th^ infestation of Am nymphs compared to control mice (Figure 1B). These results align with the trend observed in subcutaneous sensitization of mice injected with Aa TSGE (Chaudhary et al., 2021). Though we observed that α-gal specific IgG1 increased gradually during repeated sensitization, the level of α-gal specific IgE during tick feeding did not follow the same increasing pattern with *Am. americanum* repeated nymph sensitization. Low levels of sIgE were detected in a few Aa-sensitized mice as well as in Am-sensitized mice. Previous studies reported a correlation between elevated α-gal specific IgG1 and high IgE titer in AGS patients (Joral et al., 2022). Despite elevated levels of α-gal IgG1, high titer α-gal IgE was not observed in Aa nymph-sensitized mice. The most likely explanation is that tick salivary factors inoculated at the bite site inhibited the alpha-gal-sIgE-driven inflammatory response in the later infestations and aided in tick feeding (Kotal et al., 2015). Mice sensitization results imply tick bites of Aa can have low anti-alpha-gal IgE response and consistently elevated levels of alpha-gal IgG1 response. On the other hand, Am sensitization generated a low level of α-gal the IgE response, and α-gal the IgG titer remained extremely low. These trends suggest that the elevation in the titer of α-gal specific IgE following tick bites may be attributed to antibody switching induced by tick salivary prostaglandins, a phenomenon believed to play a role in driving IgE class switching and IgE production (Cabezas-Cruz et al., 2017). Since α-gal alone does not induce an IgE response (Chandrasekhar et al., 2020), there must be sensitization driven by antigen and tick salivary factors such as prostaglandin E_2_ (PGE_2_) stimulating the synthesis of the α-gal IgE antibodies (Carvallo-costa et al., 2015). Thus, an “atypical” Th2-like host response may occur during tick sensitization to produce α-gal IgE. In addition, another possible mechanism is the presence of α-gal bound lipids in the tick saliva, which could potentially trigger the release of Th2 biasing cytokines from NKT cells. As reported earlier, this immune response skewing towards Th2 could contribute to the production of α-gal IgE antibodies (Kaer and Wu 2018).

In this study, we presented evidence that alpha-gal-specific immune response of the host triggered by tick bite-sensitized mice led to an allergic reaction upon challenge with pork kidney meat. This indicates that α-gal specific sensitization of the host is caused or boosted by a particular tick expressing α-gal in saliva during a tick bite. It is also important to note that the dose of the allergen can also vary during repeat tick sensitization as nymphs are in an undifferentiated developmental stage, and lightweight engorged nymphs molt into male adults after feeding on the host (Karim et al., 2011).

We investigated host response analysis using a Nanostring approach to understand how Aa tick bite shapes AGS development in a murine model. Infestation of AGKO mice with ticks resulted in an initial burst of proinflammatory response characterized by a robust increase in the transcript of *IL-1*β as well as components of the NLRP3 inflammasome complex that is essential for IL-1β activation (Jo et al., 2016). Among inflammasome receptors, NLRP3 is unique because it is activated by diverse pathogen-associated molecular patterns (PAMP) and damage-associated molecular patterns (DAMP) from dying or injured cells. It is intriguing to speculate that an increase in *HK3* could also activate the inflammasome in a PAMP-independent manner (Wolf et al., 2016). In sum, tick infestation in mice resulted in a massive induction of pro-IL-1β by both NF-κB and MAPK pathways and its cleavage to an active form can occur by the NLPR3 inflammasome, which was also markedly upregulated in skin biopsy samples from tick bitten mice.

The IL-6 family of cytokines plays an important role in antimicrobial and antiviral immunity and provides tissue protection from infection-related injury (Tanaka et al., 2014). These cytokines often control the recruitment, adhesion, survival and effector activities of neutrophils, tissue-resident and inflammatory monocytes, and innate lymphoid cell populations including NK cells. We observed robust upregulation of both *Osm* and *Il6* and several of their downstream targets, specifically neutrophil-activating chemokines *Cxcl1* and *Cxcl5*, adhesion molecules *Icam1* and *Vcam1* (Heinze et al., 2012). While these molecules facilitate the effector function of neutrophils and the acute phase response, they also serve as lymphokines to promote the differentiation of Th1 or Th2 cells (Dienz et al., 2009, Jones and Jenkins 2018). We further observed changes in gene signatures involved in suppressing inflammation and inducing immune tolerance. While no significant change was observed in *Tbx21*, critical to Th1 differentiation in tick-infested mice, the transcription factor *Foxp3* was significantly induced following the 2nd infestation, consistent with a Treg response to suppress inflammation (Ono et al., 2007). We also observed increased transcription of *Havcr2*, which is expressed on Treg cells and could inhibit both Th1 and Th17 responses (Sanchez-Fueyo et al., 2003, Gautron et al., 2014).

Repeated exposure to tick saliva has been suggested to skew polarization of the immune response toward a Th2 profile, leading to the development of allergies and suppression of proinflammatory response. In this context, we observed increased transcripts of Th2 cytokines *Il33* and *Il4*, as well as their receptors, following subsequent tick infestations that were also associated with increased production of IgG and IgE measured by ELISA (Coffman et al., 1986, Schmitz et al., 2005) (**Figure 6**). Further, we also found evidence of tick bite induced robust *Tlr1*, *Tlr2*, *Tlr6*, and *Tlr8* signaling and *MyD88* induction, both of which could contribute to IgE production as suggested by Chandrasekhar and colleagues (2019). An increase in *Alox15*, *Atf4* and *Xbp1* transcripts which modulate ER stress response for efficient immunoglobulins secretion was further noted (So, 2018).

Prostaglandin E2 (PGE_2_) is one of the most abundant bioactive molecules in tick saliva and has been implicated in driving IgE class switching and the production of IgE (Cabezas-Cruz et al., 2017). We observed upregulation of *Ptgs2* transcription, a key enzyme in prostaglandin biosynthesis, as well as prostaglandin E receptor *Ptger4* and bradykinin receptor *Bdkrb1* (Kosaka et al., 1994). Bradykinin can lead to the release of prostaglandins (Jose et al., 1981). It is tempting to speculate that the upregulation of *Ptger4* and *Bdkrb1,* and the production of PGE_2_, might have contributed to class switching from existing B cell clones producing anti-alpha-Gal IgM and/or IgG to anti-alpha-Gal IgE-producing B cells.

IL-20 subfamily of cytokines protects epithelial cells against extracellular pathogens and plays a vital role in wound healing and homeostasis of the tissue epithelial layer (Rutz et al., 2014). IL-19 and IL-24 are produced primarily by myeloid cells principally through TLR activation, although epithelial cells and Th2 cells can also generate both cytokines under certain conditions (Ouyang et al., 2011). We observed increased transcription of *Il19* and *Il24* following tick bites, which peaked following the 3^rd^ infestation. Lipocalin 2 (*Lcn2*), which is important in controlling bacterial infection in mice (Wang et al., 2019) and could be induced by the IL-20 subfamily was upregulated (Behnsen et al., 2014). IL-24 suppresses IL-1β expression in keratinocytes to dampen inflammatory responses and stimulate keratinocytes for tissue repair (Myles et al., 2013). Further, it induces factors that play critical roles in modulating inflammatory responses, enhancing granulation tissue formation, and inducing angiogenesis.

The complement cascade offers the first line of defense against bites from ectoparasitic ticks and has been shown to play a part in acquired tick resistance (ATR) (Kitsou et al., 2021). Following infestation of AGKO mice with *Aa*, complement component 3 (C3) and complement receptors *C3ar1* and *C5ar1* were upregulated while Cd59a, a potent complement membrane attack system inhibitor, was downregulated. We observed upregulation of *Plau* and *Plaur* which could cause activation of plasminogen into plasmin (PL), a broad-spectrum serine-protease. Interestingly, plasmin could initiate a classical complement cascade resulting in complement activation and might contribute to the recruitment of basophils as observed in tick attachment sites in guinea pig models Allen et al., 1979). We observed upregulation of several genes such as *Il3ra*, *Hdc*, *Lyn*, *Syk*, *Fcer1a* and *Pik3cd* which prime mature basophils and mast cells resulting in degranulation following IgE binding or exposure to anaphylatoxins and therefore might also contribute to ATR (Yoshikawa et al., 2021).

## Conclusion

Here we describe key immune determinants linked to host response and AGS induction following *Am. americanum* repeated tick bite using AGTKO mice model. We also demonstrated and established a method that included the use of nymph to investigate the role of ticks in the induction of AGS—following tick bite. We also reported that presence of α-gal antigens in tick plays a critical role in the sensitization of host against α-gal following tick bite. This nanostring dataset will be important for understanding critical host pathways, immune gene linked to AGS induction in host following tick bite. Our study also validated the fact that high α-gal IgG1 titer is an indicator of AGS development in the host and tick bite plays role in boosting host response against α-gal antigen leading to sensitization. Our findings also demonstrated that presence of α-gal in tick is critical factor for sensitization against α-gal in host during tick bite.

## Ethics Statement

All animal experiments were conducted per the guidelines in the Guide for the Care and Use of Laboratory Animals of the National Institutes of Health, USA. The Institutional Animal Care and Use Committee of the University of Southern Mississippi approved the protocol for the tick blood-feeding on sheep.

## Author Contributions

Conceptualization: Shahid Karim

Methodology: Surendra Raj Sharma, Shailesh Chaudhary, Scott Commins, Shahid Karim

Data curation: Surendra Raj Sharma, Shailesh Chaudhary, Julia Vorobiov, Scott Commins, Shahid Karim

Funding acquisition: Shahid Karim

Investigation: Surendra Sharma, Shailesh Chaudhary, Julia Vorobiov, Scott Commins, Shahid Karim

Project administration: Shahid Karim

Resources: Scott Commins, Shahid Karim

Supervision; Shahid Karim

Validation: Surendra Raj Sharma, Scott Commins, Shahid Karim

Writing, original draft: Surendra Raj Sharma, Shailesh Chaudhary, Scott Commins, Shahid Karim

Writing, review & editing: Surendra Sharma, Shailesh Chaudhary, Scott Commins, Shahid Karim

All authors read and approved the manuscript.

